# A lifespan-scale single-cell atlas defines an early-childhood immunometabolic transition linked to age-referenced immune states

**DOI:** 10.64898/2026.09.11.750312

**Authors:** Jiayi Zheng, Yu Shi, Yichen Sun, Yalin Wang, Ruoxin Li, Peng Wang, Shiru Chang, Lanxin Pan, Huiling Tan, Tong Yue, Kun Qu, Xueying Zheng, Chuang Guo, Jianping Weng

## Abstract

Childhood represents a critical period with rapid metabolic and immune remodeling, during which perturbations may have lasting consequences for health. We integrated single-cell RNA sequencing (scRNA-seq) profiles of peripheral blood mononuclear cells (PBMCs) from 2,008 healthy individuals aged 0–100 years. Age-resolved modeling highlighted an early-childhood transition centered at age 6, characterized by coordinated remodeling of metabolism-related transcriptional programs, particularly in monocyte and B-lineage cells. Matched serum metabolomics and lipidomics in a pediatric subcohort detected a temporally concordant wave of metabolite and lipid changes around ages 6–8 and extensive metabolite/lipid–transcript associations, prominently involving steroid and phospholipid species. Building on these findings, we developed an immunometabolic transition score and age-specific reference chart, which were evaluated in an independent healthy cohort and applied to quantify deviations in pediatric immune disorders. These findings highlight a coordinated immunometabolic transition in early childhood and provide an age-referenced framework for distinguishing atypical immune-state variation from healthy age-associated variation.

---

The immune system undergoes rapid and essential maturation in early life^1^, resulting in marked differences between neonatal and adult peripheral immune-cell composition and functional states^1,2^. During this period, immunometabolism, an important determinant of immune cell maturation and function^3,4^, is increasingly recognized as a contributor to immune health, as disruption of metabolic signals in childhood has been linked to immune-mediated disease onset and chronic inflammatory states^5,6^. Therefore, characterizing healthy immune remodeling across childhood in relation to systemic metabolic changes may clarify how age-associated immune states are established and how they are altered in disease.

Recent advances in single-cell RNA sequencing (scRNA-seq) have enabled the characterization of age-associated immune variation at cell-type resolution in circulating immune cells^7–9^. However, existing PBMC single-cell data often have sparse and uneven representation of pediatric ages, limiting the characterization of immune remodeling across childhood. Moreover, these datasets have rarely been integrated with complementary molecular layers, such as metabolomics and lipidomics, to capture coordinated systemic changes. Methodologically, coarse age binning can dilute age-localized signals, while multi-omics studies increasingly support temporally clustered, nonlinear remodeling across molecular layers^10,11^. In addition, analyses that rely on discrete comparisons between children and adults may conflate features specific to immune development and aging. Together, these limitations leave the timing, pattern, and systemic metabolic context of childhood immune remodeling incompletely resolved within the human lifespan.

Age-associated immune features have been explored as biomarkers of immune health^7,8^. Given the dynamic physiological changes during childhood, interpreting these features in pediatric populations requires accounting for normal developmental variation^12^. In pediatric practice, growth references and charts are widely used in traits such as height, weight, and brain volume to identify atypical deviations from healthy age-matched reference populations^13,14^. However, comparable age-referenced approaches for interpreting individual immune-state variation remain underdeveloped.

Here, we integrated PBMC single-cell transcriptomes from 2,008 healthy individuals spanning 0–100 years with independent cohorts for validation. Using age-resolved modeling, we revealed age-localized transitions of intensified transcriptional reprogramming and highlighted a prominent early-childhood transition enriched for metabolism-related transcriptional programs. We then placed this transition in a systemic context using serum metabolomic and lipidomic profiling in a pediatric subcohort. Finally, leveraging genes defining the early-childhood immunometabolic transition, we developed a transition score together with an age-specific reference chart to quantify individual deviations in pediatric immune disorders.

## Results

### Single-cell atlas of peripheral immune cells across the human lifespan

To characterize age-associated remodeling of peripheral immune cells, we compiled single-cell RNA-sequencing (scRNA-seq) profiles of peripheral blood mononuclear cells (PBMCs) from 2,008 healthy individuals spanning 0–100 years (46.81% male, Fig. 1A). This discovery dataset included 1,952 donors aggregated from 28 public studies (Supplementary Table 1; median age 55 years, interquartile range [IQR], 37–69). To improve representation of pediatric ages that are sparse in public resources, we additionally generated PBMC scRNA-seq data from 56 healthy donors (median age 10.86 years, IQR, 7.75–14.00) (Supplementary Table 2). After doublet removal and quality control filtering, the discovery atlas comprised 5,749,312 cells. Following batch correction and dimensionality reduction (Extended Data Fig. 1A), we annotated cell-type labels using CellTypist^15^ and identified 26 immune cell subsets (Fig. 1B), which were further supported by canonical marker gene expression (Extended Data Fig. 1B).

**Fig. 1:**
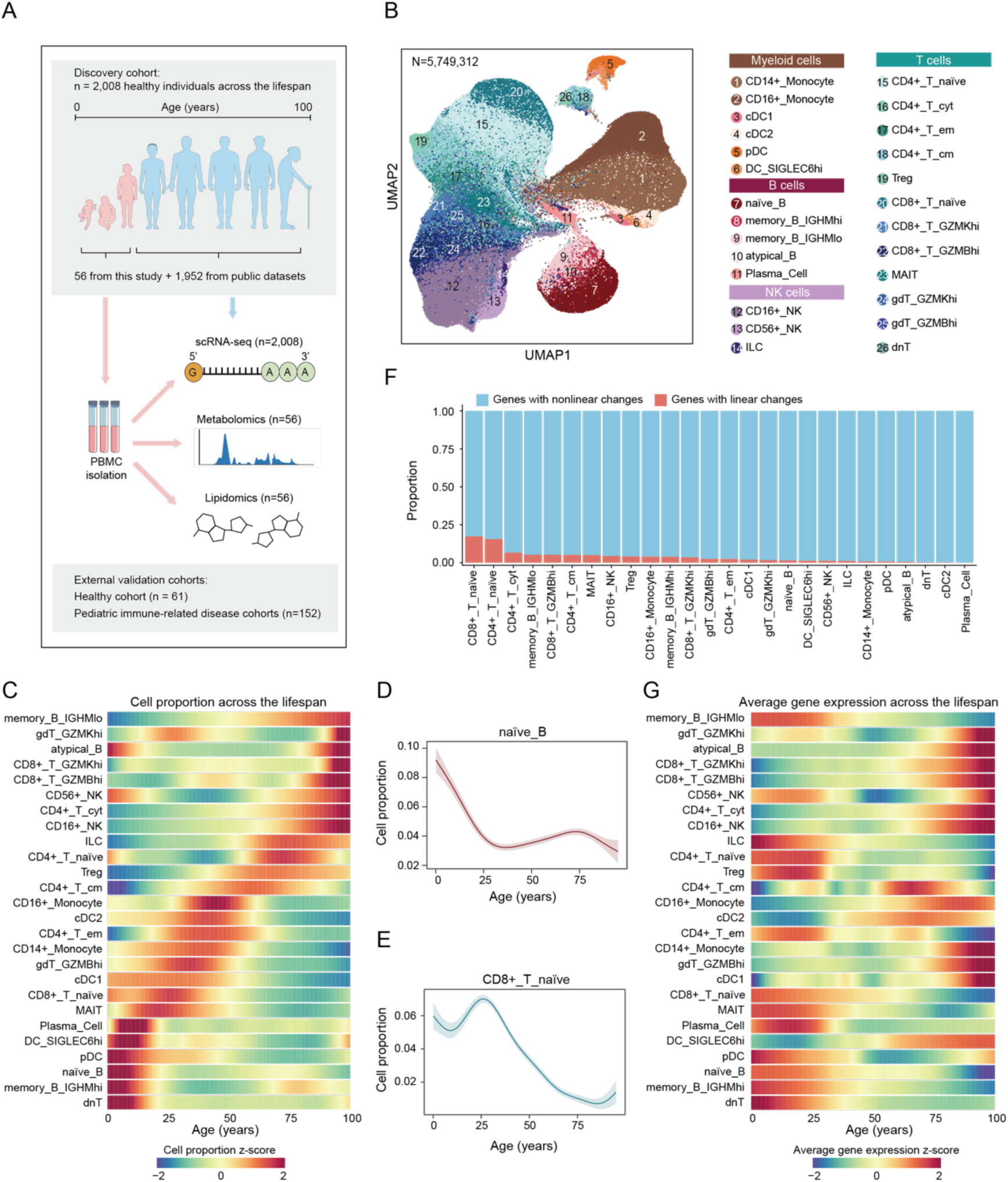
Study design and single-cell transcriptomic atlas of human peripheral immune cells across the lifespan. **A,** Overview of the study design and data. **B,** Uniform manifold approximation and projection (UMAP) visualization of 5,749,312 PBMCs from the discovery cohort, colored by 26 annotated immune cell populations. **C,** Heatmap showing z-scored proportions of immune cell subsets across the human lifespan. **D-E,** Representative age-associated trajectories of immune cell subset proportions across the human lifespan. Lines represent fitted smooth trajectories, and shaded areas indicate 95% confidence intervals. **F,** Stacked bar plot showing the proportion of genes showing nonlinear versus linear changes with age across different immune cell types. **G,** Heatmap showing aggregate transcriptional profiles of immune cell subsets across the lifespan.

We next quantified age-associated changes in PBMC composition. For each donor, we calculated the proportion of each immune subset and modeled donor-level proportions as smooth functions of chronological age (Methods). This analysis revealed heterogeneous, life-stage-dependent patterns across immune subsets (Fig. 1C). During childhood, several immune cell subsets showed relatively higher proportions, including naïve B cells and CD8⁺ naïve T cells, which subsequently declined with age (Fig. 1D, E). In contrast, GZMK^hi^ CD8⁺ T cells and cytotoxic CD4⁺ T cells showed relatively lower proportions during childhood compared with later ages (Extended Data Fig. 1C, D). Several cell types showed dynamic patterns that differed between childhood and adulthood, such as MAIT, which increased in proportion during childhood but predominantly declined across adulthood and older ages (Extended Data Fig. 1E).

We then characterized the temporal dynamics of age-associated transcriptional remodeling within immune cell subsets using linear and nonlinear modeling approaches (Methods). Linear association analyses identified relatively small sets of age-associated genes, with the largest numbers detected in naïve CD4^+^ and CD8^+^ T cells (Extended Data Fig. 2A). In contrast, models allowing for nonlinear relationships identified a substantially larger number of age-associated genes across immune cell subsets (Methods, Extended Data Fig. 2B). Overall, 93.94% of age-associated genes showed age associations captured by nonlinear models across the lifespan (Extended Data Fig. 2C), with certain cell types, such as plasma B cells and cDC2, approaching 100% (Fig. 1F). Accordingly, the nonlinear framework detected an average of 1,864 age-associated genes per immune cell subset compared with 120 from the linear analysis (Supplementary Table 3). We further visualized overall transcriptional trajectories of immune cells by averaging normalized expression across all genes. These aggregate transcriptional profiles also exhibited life-stage-dependent patterns across the lifespan (Fig. 1G). Notably, the elevated transcriptional states of naïve B cells and naïve CD8⁺ T cells during childhood coincided with similar age-related patterns in cellular proportions (Fig. 1C, G). Together, these analyses indicate that age-associated transcriptional remodeling is predominantly nonlinear.

### Quantification of transcriptional changes across the lifespan uncovers waves of age-associated genes

The predominance of nonlinear age-associated immune remodeling indicates that transcriptional changes are unevenly distributed across the lifespan. To reveal when these remodeling events are most concentrated, we therefore applied differential expression-sliding window analysis (DE-SWAN)^10^ to localize and quantify age-localized transcriptional changes across immune cell subsets (Methods). This analysis revealed wave-like patterns of differentially expressed genes (DEGs) across age, and the timing of these waves varied across immune cell subsets (Fig. 2A-D). T cell subsets exhibited the most frequent age-localized waves across the lifespan (Fig. 2A). By contrast, several immune cell subsets displayed more restricted and age-specific waves, such as atypical B cells in early childhood (centered at 6 years) (Fig. 2C). These patterns suggest that distinct immune compartments contribute to transcriptional remodeling at different stages of life.

**Fig. 2:**
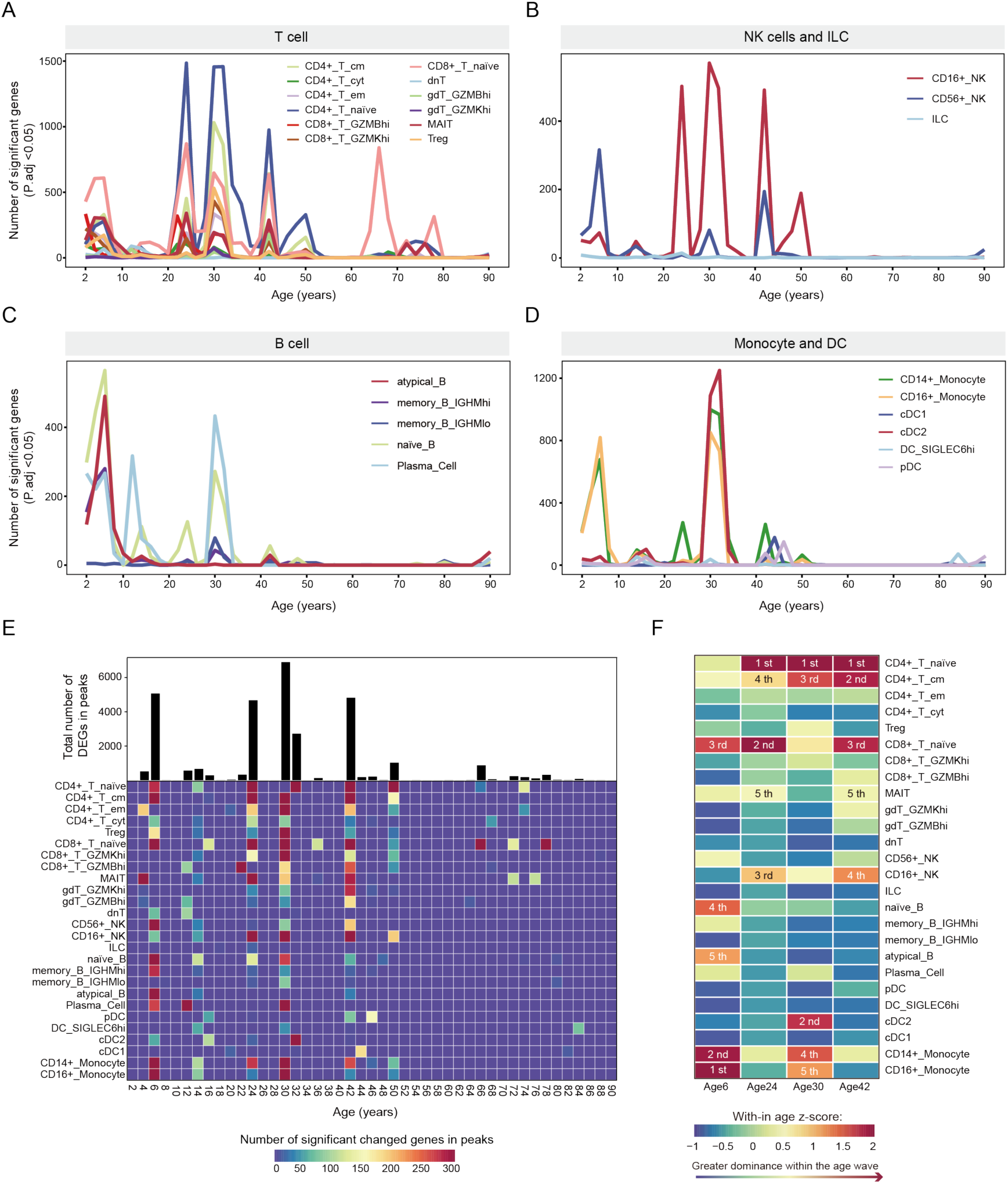
Waves of age-associated gene expression across immune cell types. **A–D,** Number of age-associated differentially expressed genes (DEGs) identified within each age window in T cells (A), NK cells and ILCs (B), B cells (C), and monocytes and dendritic cells (D). Significant DEGs were defined as adjusted P value < 0.05. **E,** Heatmap summarizing the number of DEGs at each peak across all cell types. Color intensity indicates the number of DEGs contributing to each peak within each cell type. The bar plot above shows the total number of DEGs summed across cell types at each peak age center. **F,** Heatmap showing z-scored DEG counts for each cell type within the major transition windows centered at 6, 24, 30, and 42 years. Higher values indicate a stronger relative contribution of transcriptional changes within the corresponding transition window. Ranked labels denote the top five contributing cell types at each age window.

To identify transition windows with the most substantial remodeling, we quantified the number of DEGs across age windows for each immune cell subset and identified local peaks in the DEG waves (Methods). Age windows centered at 6, 24, 30, and 42 years emerged as major periods of transcriptional remodeling, with broadly distributed and pronounced DEG peaks across immune cell populations (Fig. 2E, Supplementary Table 4). The age windows centered at 24, 30, and 42 years align with peaks of remodeling reported in previous proteomic and bulk transcriptomic studies^10,11,16^. Beyond these reported transition periods in adulthood, we also highlight an early-childhood window centered at 6 years.

We next assessed cell-type contributions to each age-localized window by ranking immune cell subsets according to the number of DEGs in each transition window (Fig. 2F). This analysis revealed a stage-dependent shift in the dominant contributors. Compared with adult transition windows, the transition centered around 6 years was characterized by substantially greater contributions from the monocyte and B cell compartments. Notably, naïve B cells and atypical B cells exhibited their highest contribution to transcriptional remodeling across the lifespan within this transition window (Fig. 2C, Fig. 2F). In contrast, some CD4⁺ T cell subsets, including naïve and central memory CD4⁺ T cells, although displaying local peaks around 6 years, contributed less prominently than in adult transition windows, where they DEG burdens ranked among the highest (Fig. 2A, Fig. 2F). CD8⁺ naïve T cells, meanwhile, remained prominent contributors to transcriptional remodeling in both childhood and adult transitions (Fig. 2F). Together, these results delineate discrete age windows of intensified transcriptomic remodeling and highlight distinct cell type-specific DEG patterns during early childhood compared with adulthood.

### The 6-year transition is marked by transcriptional remodeling of metabolic pathways

Following the stage-dependent shift in dominant contributing immune subsets across the windows centered at 6, 24, 30, and 42 years (Fig. 2F), we next asked whether these windows were also associated with distinct functional programs. Most (91.28%) window-associated DEGs were detected in only one age-localized window, including 5,277 unique to the 6-year transition, suggesting that these windows capture distinct transcriptional programs (Extended Data Fig. 3A). Gene Ontology (GO) enrichment analysis of DEGs detected within these transition windows revealed that the transition centered at 6 years showed a prominent metabolic signature, with preferential enrichment of carbohydrate metabolism and small-molecule metabolic pathways (Fig. 3A). This pattern was particularly pronounced in monocyte and B-cell lineage compartments, where metabolism-related terms dominated the top enriched pathways (Supplementary Table 5). By comparison, DEGs from the 24-, 30- and 42-year transitions were more consistently enriched for programs related to protein homeostasis, cell activation and kinase-mediated signaling, and responses to extracellular stimuli, respectively (Extended Data Fig. 3B–D; Supplementary Table 6–8).

**Fig. 3:**
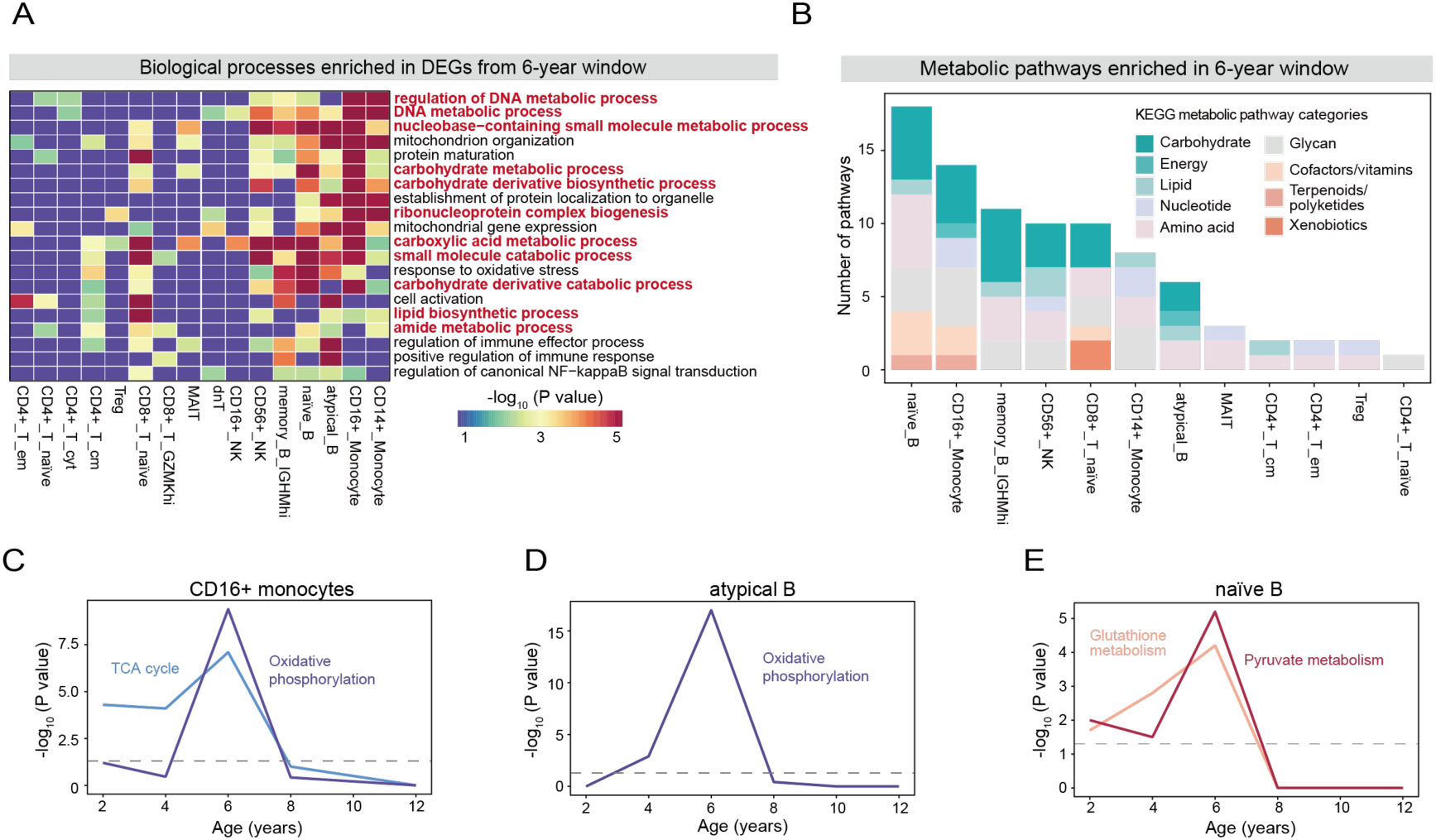
The 6-year transition is marked by transcriptional remodeling of metabolic pathways. **A,** Heatmap showing the enrichment of biological processes among DEGs in the 6-year transition window across immune cell subsets. Metabolism-related terms are highlighted in red. **B,** Stacked bar plot showing the number of enriched metabolic pathways in the 6-year transition window across immune cell subsets. Colors indicate Kyoto Encyclopedia of Genes and Genomes (KEGG) metabolic pathway categories. **C–E,** Line plots illustrating the significance of metabolic pathway enrichment, expressed as −log_10_(P value), for (C) TCA cycle and oxidative phosphorylation in CD16^+^ monocytes, (D) oxidative phosphorylation in atypical B cells, and (E) glutathione and pyruvate metabolism in naïve B cells. Dashed grey lines indicate P = 0.05.

To characterize metabolic features of the 6-year transition window in greater detail, we curated metabolic pathways from the Kyoto Encyclopedia of Genes and Genomes (KEGG) database and tested whether DEGs in the 6-year transition were overrepresented in these pathways (Methods). KEGG metabolic pathways were grouped into 10 major categories (Supplementary Table 9), and nine of the ten categories contained pathways that were significantly overrepresented among DEGs from the 6-year window (Extended Data Fig. 3E). Carbohydrate metabolism accounted for the largest fraction of significantly enriched pathways. At the cell-type level, 12 immune cell subsets exhibited significant enrichment of KEGG metabolic pathways among their 6-year window-associated DEGs, with naïve B cells showing the largest number of enriched metabolic pathways, followed by CD16⁺ monocytes (Fig. 3B), consistent with their prominent transcriptional changes at this window (Fig. 2F).

We next examined the temporal dynamics of metabolism-related pathway enrichment across childhood. The number of significantly enriched metabolic pathways and metabolism-related genes increased progressively in early childhood, reaching a peak in the 6-year window (Extended Data Fig. 4A). Across KEGG metabolic categories, enrichment generally peaked around 6 years, with pathways related to carbohydrate metabolism accounting for the largest fraction of overrepresented genes during the 2–6-year interval (Extended Data Fig. 4B).

The peak centered at 6 years was also evident at the level of specific metabolic pathways within individual immune cell subsets. In CD16⁺ monocytes, genes annotated to the tricarboxylic acid (TCA) cycle and oxidative phosphorylation were significantly overrepresented among window-associated DEGs across the 2–6-year interval, with the strongest enrichment observed in the 6-year window (Fig. 3C), consistent with previously reported changes in carbon and energy metabolism in early childhood^2^. A similar pattern for oxidative phosphorylation was also observed in atypical B cells, indicating that oxidative phosphorylation–related transcriptional remodeling is not restricted to the myeloid compartment (Fig. 3D). We also observed dynamic enrichment of other metabolism-related pathways. For example, glutathione and pyruvate metabolism showed pronounced enrichment dynamics in naïve B cells and IGHM^hi^ memory B cells, with the strongest signals at the 6-year window (Fig. 3E, Extended Data Fig. 4C). Together, these results suggest dynamic remodeling of metabolic gene programs across early childhood, with pathway-level signals most concentrated around 6 years.

### A wave of age-related serum metabolites and lipids in early childhood

To evaluate whether the early-childhood remodeling of metabolism-associated transcriptional programs is accompanied by systemic molecular changes, we profiled serum metabolites in a pediatric subset of the lifespan cohort (n = 56, ages 2–21 years, Supplementary Table 2) using untargeted metabolomics. DE-SWAN identified an age-localized wave of differential metabolite abundance within the age window spanning 6–8 years (Fig. 4A). Lipids and lipid-like molecules represented the largest metabolite superclass among differentially abundant metabolites within the 6–8-year wave, with 56 species increased and 32 decreased relative to adjacent age windows (Fig. 4B). Within the lipids and lipid-like molecules, fatty acyls were the most represented class, followed by steroids and steroid derivatives (Fig. 4B).

**Fig. 4:**
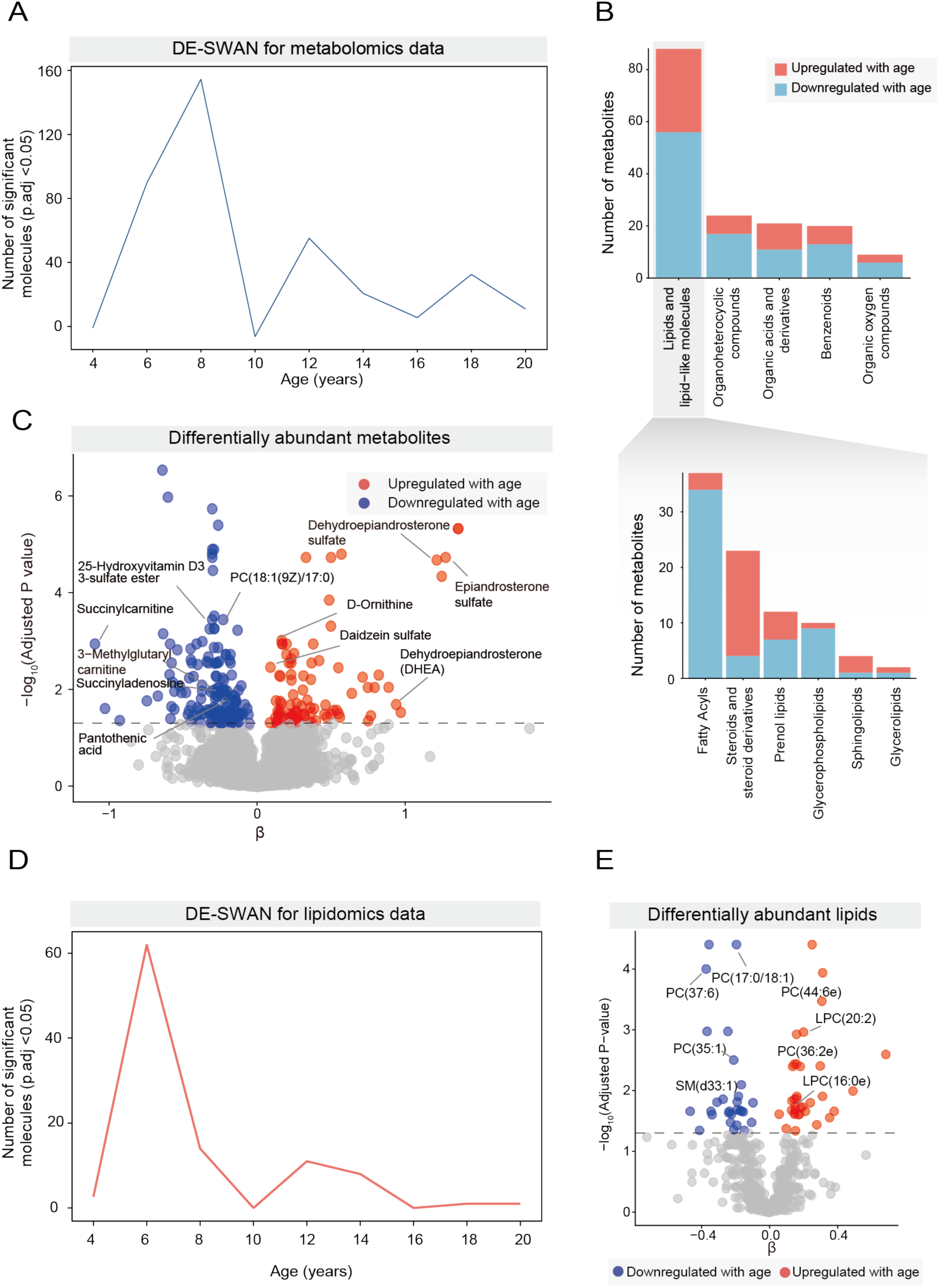
Serum metabolomic and lipidomic waves in early childhood. **A,** The number of differentially abundant metabolites identified from the metabolomic data by DE-SWAN across sliding-window age centers. **B,** Numbers of differentially abundant metabolite species across metabolite classes. The upper panel shows the five most represented metabolite superclasses. The lower panel further shows the five most represented classes within the Lipids and lipid-like molecules superclass. Stacked bars indicate metabolites that were upregulated (red) or downregulated (blue) with age. **C,** Volcano plot showing differentially abundant metabolites in the 6–8-year windows. Blue and red points indicate metabolite species decreased and increased with age, respectively. Dashed grey lines indicate P = 0.05. **D,** Number of differentially abundant lipids from the lipidomic data identified by DE-SWAN across sliding-window age centers. **E,** Volcano plot showing differentially abundant lipids in the 6-year window. Blue and red points indicate lipid species decreased and increased with age, respectively. Dashed grey lines indicate P = 0.05.

Within this metabolic wave, significant age-related changes in abundance were observed in metabolites associated with carbohydrate, nucleotide, and lipid metabolism. For example, succinylcarnitine (linked to TCA cycle intermediates) and succinyladenosine (derived from purine metabolism) decreased in abundance relative to adjacent age windows (Fig. 4C). In contrast, multiple steroids and their derivatives, including dehydroepiandrosterone (DHEA) and dehydroepiandrosterone sulfate (DHEAS), increased in abundance within the 6–8-year window (Fig. 4C).

Given the prominence of lipid-related metabolites in the 6–8-year metabolite wave, we further performed untargeted lipidomics in the same cohort. Lipidomics similarly revealed an age-localized wave of significantly changing lipid species around age 6 (Fig. 4D), dominated by an age-related increase in polyunsaturated lipids (PU; Extended Data Fig. 5A). Among lipid classes, phosphatidylcholines (PCs) exhibited the largest number of changing species (Extended Data Fig. 5B), with both increasing (e.g., PC(44:6e)) and decreasing (e.g., PC(35:1)) species (Fig. 4E). Together, these results define an early-childhood window of metabolomic and lipidomic remodeling.

### Metabolite– and lipid–transcript associations during the early-childhood transition window

We next assessed whether circulating molecules that change within the early childhood transition covary with immune transcriptional changes over the same interval. Spearman association analysis identified 20,281 significant metabolite–transcript associations across 22 immune cell types (adjusted P value <0.05, |ρ| > 0.2, Fig. 5A). Steroids and steroid derivatives accounted for the largest number of associations (n = 3,860), followed by carboxylic acids and derivatives (n = 1,822) (Fig. 5B). Steroid- and steroid derivative-associated transcripts showed cell-type-dependent functional enrichment patterns (Fig. 5C, Supplementary Table 10). For example, in CD16⁺ monocytes, steroid- and steroid derivative-associated transcripts were enriched for pathways related to cytokine production (Fig. 5C), whereas in central memory CD4^+^ T cells, steroid- and steroid derivative-associated transcripts were enriched for regulation of T cell differentiation. Carboxylic acid and derivative-associated transcripts were enriched for pathways including carbohydrate metabolism (e.g., Hexose and monosaccharide metabolic process in IGHM^hi^ memory B cells) and regulation of leukocyte activation in MAIT (Fig. 5D, Supplementary Table 11).

**Fig. 5:**
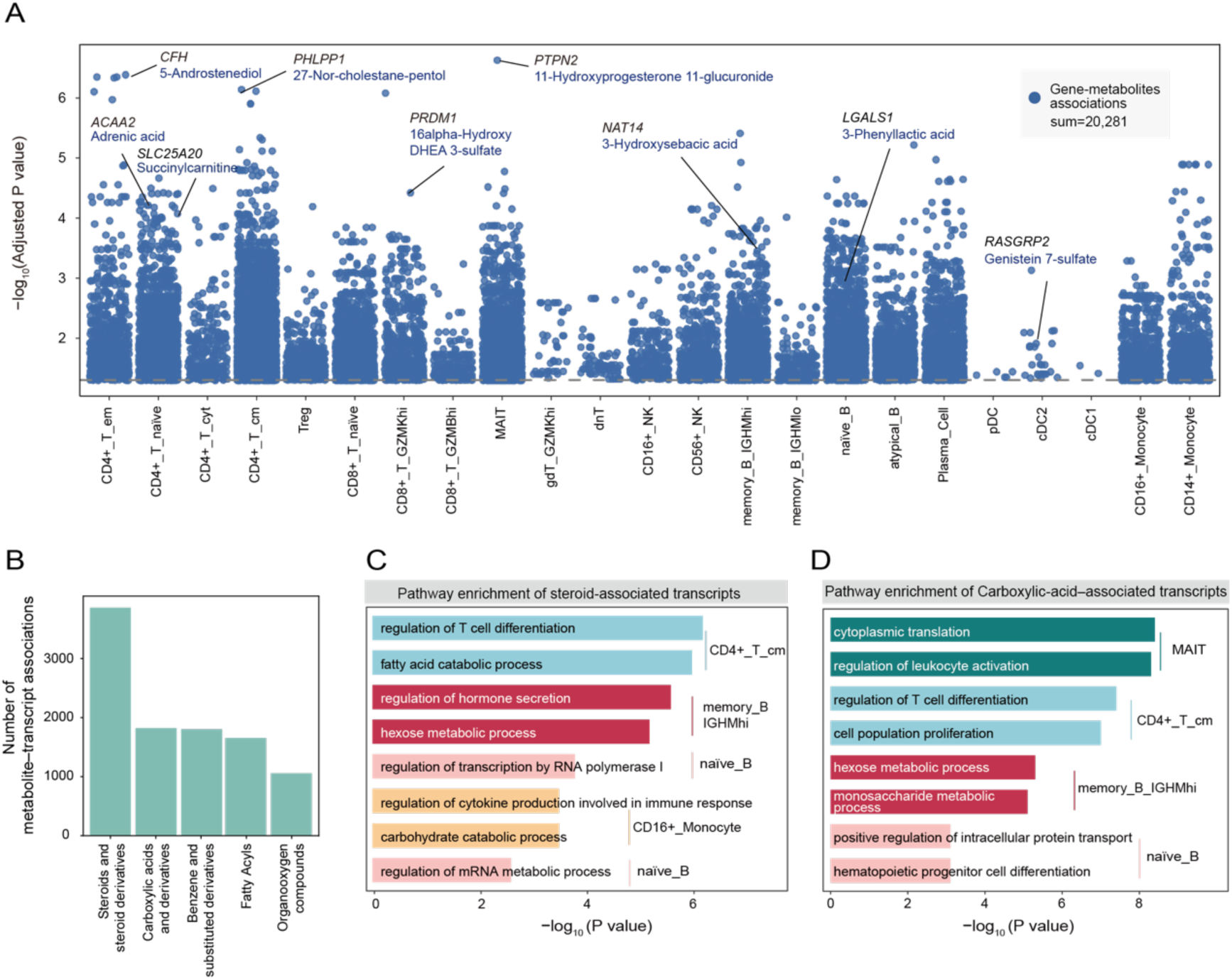
Serum metabolomic waves around age 6 are associated with immune transcriptional remodeling. **A,** Dot plot showing the significant metabolite–transcript associations across immune cell subsets. Dashed grey lines indicate P = 0.05. **B,** Bar plot showing metabolite classes with more than 1,000 significant metabolite–transcript associations. **C,** Bar plots showing the functional enrichment of steroid- and steroid derivative-associated transcripts. **D,** Bar plots showing the functional enrichment of carboxylic acid and derivative-associated transcripts.

We then linked lipid abundances to expression of window-associated DEGs within the 6-year wave and identified 5,856 significant lipid–transcript associations (adjusted P value <0.05, |ρ| > 0.2, Extended Data Fig. 6A). PCs (n = 2,450) and lysophosphatidylcholines (LPCs; n = 924) accounted for the largest numbers of associations (Extended Data Fig. 6B). PC-associated transcripts were enriched for pathways related to immune co-stimulation and intracellular signaling (Extended Data Fig. 6C, Supplementary Table 12). LPC-associated transcripts showed enrichment for pathways involved in molecular processing and post-transcriptional regulation (Extended Data Fig. 6D, Supplementary Table 13).

At the molecular level, the integrative association analysis identified metabolite/lipid–transcript pairs with potential biological relevance (Fig. 5A, Extended Data Fig. 6A). Several of the significant correlations linked metabolism-related genes with circulating metabolites. In naïve CD4⁺ T cells, the expression of *SLC25A20* (encoding the carnitine–acylcarnitine translocase) showed a positive association with serum succinylcarnitine abundance (adjusted P value = 9.80 × 10⁻^5^, ρ = 0.59, Fig. 5A). These associations also linked circulating molecules to immune-regulatory transcriptional programs. For example, LPC species have been implicated in autoimmune pathogenesis^17^. In naïve B cells, LPC (20:2) abundance was associated with *ARID3A* expression (adjusted P value = 2.40 × 10⁻^5^, ρ = −0.62, Extended Data Fig. 6A), a transcription factor whose hyperactivity during early B-lineage development has been linked to autoreactivity of mature B cells^18^. Together, these analyses connect the serum metabolomic and lipidomic wave around ages 6-8 years with cell type-resolved transcriptional remodeling in the corresponding early-childhood age window, highlighting circulating steroids and phospholipids as major correlates of this temporally concordant immunometabolic transition.

### A transition score and an age-specific reference anchored to the 6-year window

Building on the identified early-childhood transition around 6 years, we developed a transition score to quantify this immunometabolic transition at the individual level (Fig. 6A). Transition-associated DEGs that significantly correlated with serum metabolite or lipid levels were used for feature selection, yielding 140 features for model construction (Supplementary Table 14, Methods). The selected features were most frequently derived from naïve CD8⁺ T cells, followed by MAIT and atypical B cells (Extended Data Fig. 7A). Permutation importance analysis identified the top 10 contributors to the scoring model, including genes involved in immune trafficking or signaling genes (*e.g.*, *SELL*, *FCER1G*) and metabolism-associated genes (*e.g.*, *LGALS1*) (Extended Data Fig. 7B). A self-paced ensemble classifier was then trained to distinguish individuals before and after the 6-year age point, generating a transition score ranging from 0 (pre-6-year transition state) to 1 (post-6-year transition state).

**Fig. 6:**
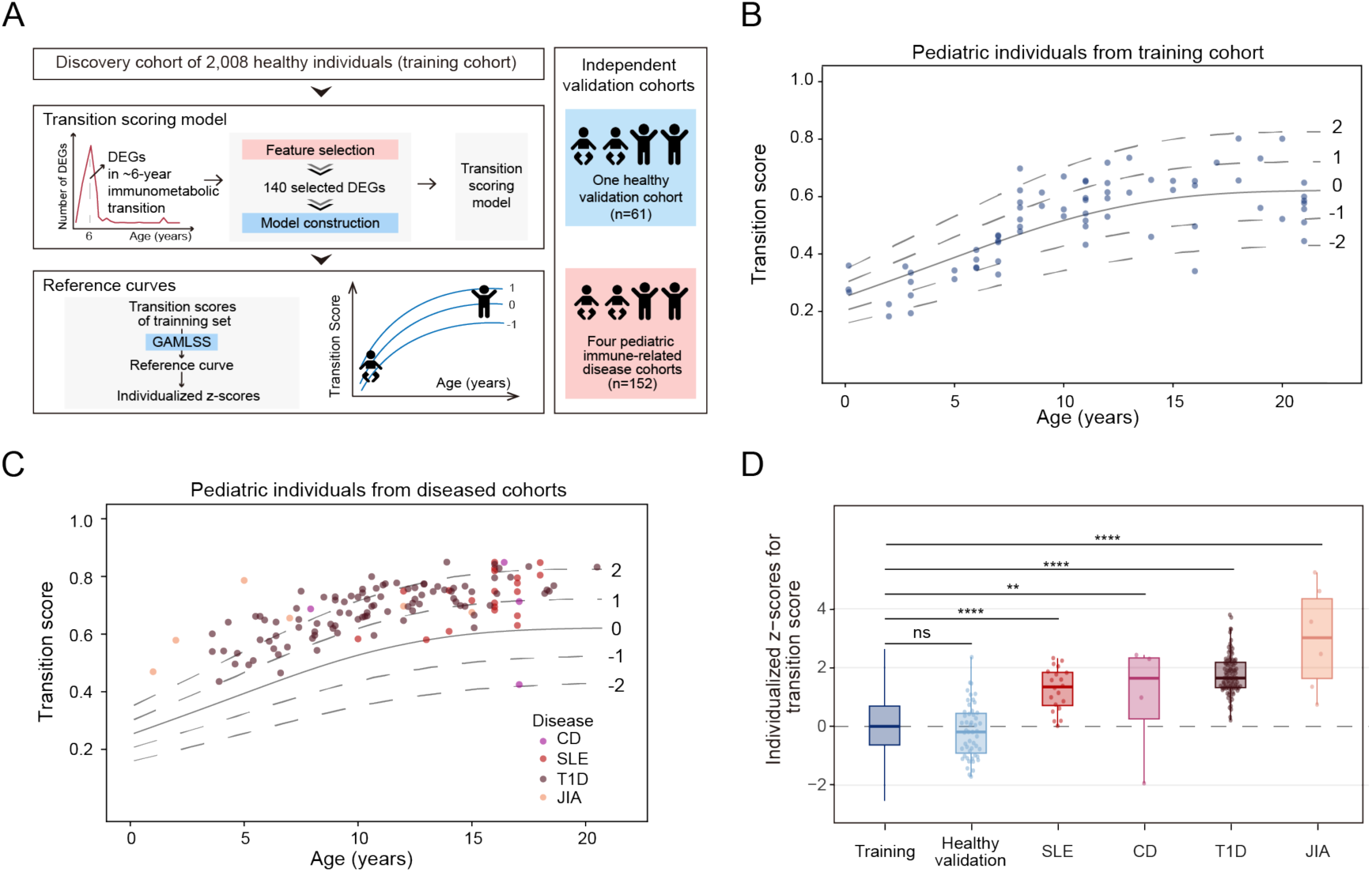
Construction of the transition score and age-specific reference chart based on the 6-year transition. **A,** Workflow for constructing the transition scoring model and age-referenced chart. **B–C,** Age-specific reference charts for the transition score in pediatric populations. The solid line represents the age-specific median of the healthy reference population (z-score = 0), and the dashed lines indicate z-scores ranging from −2 to 2. Points represent the healthy pediatric individuals (B) and the pediatric patients with immune-related diseases (C). **D,** Box plot comparing the individualized z-scores among the healthy reference population, healthy validation cohort, and pediatric immune-related disease cohorts. Statistical significance was assessed by permutation testing and an adjusted P value < 0.05 was considered statistically significant: ns, not significant; ** adjusted P < 0.01; **** adjusted P < 0.0001; SLE: systemic lupus erythematosus; CD: Crohn’s disease; T1D: type 1 diabetes; JIA: Juvenile idiopathic arthritis.

We further examined the metabolite and lipid associations of these selected features. The selected features exhibited metabolite and lipid class associations similar to those observed across all associations, with steroids and steroid derivatives remaining the predominant metabolite class (Extended Data Fig. 7C), whereas PCs remained the most frequently associated lipids (Extended Data Fig. 7D). The selected features also included associations with metabolites of known immunological relevance. For example, *LGALS1* in atypical B cells was associated with 3-phenyllactic acid (Fig. 5A), a microbiota-derived metabolite previously implicated in autoimmune disease^5,19^. These findings suggest that the transition score may capture biologically meaningful features linked to immune health.

To benchmark individual immune states in an age-aware context, we constructed an age-specific reference chart of transition scores using generalized additive models for location, scale, and shape (GAMLSS) (Methods; Extended Data Fig. 8A, B; Supplementary Table 15). The reference trajectory increased rapidly in early childhood and then stabilized into adulthood (Fig. 6B, Extended Data Fig. 8C). This reference framework enables the calculation of individualized z-scores relative to the age-specific distribution of the healthy reference population (Methods). We then evaluated the reference framework in several independent cohorts using permutation testing (Supplementary Table 16, Methods). In an independent healthy cohort, median z-scores did not significantly differ from the training reference (adjusted P value = 0.306; Extended Data Fig. 8D). In contrast, pediatric cohorts with immune-related conditions showed significant z-score deviations relative to healthy individuals, including Crohn’s disease (CD; median = 1.64, adjusted P value = 4.12×10^-3^), systemic lupus erythematosus (SLE; median = 1.34, adjusted P value < 1×10^-4^), type 1 diabetes (T1D; median = 1.64, adjusted P value < 1×10^-4^), and juvenile idiopathic arthritis (JIA; median = 3.01, adjusted P value < 1×10^-4^) (Fig. 6C, D). Together, these results provide a transition score and reference chart for quantifying deviations from typical age-associated immune trajectories in pediatric cohorts.

## Discussion

In this study, we assembled a lifespan-scale PBMC single-cell transcriptomic atlas to reveal when and in which immune compartments age-associated remodeling is most pronounced. Across immune cell subsets, age-associated transcriptional changes were predominantly nonlinear and concentrated within discrete age windows, indicating that immune remodeling is temporally structured rather than uniformly progressive. A prominent early-childhood transition around age 6 was distinguished by broad remodeling of metabolism-related transcriptional programs and accompanied by a temporally concordant serum metabolomic and lipidomic wave. Building on this transition, we further developed a transition score and age-specific reference chart to provide an age-aware framework for interpreting individual immune states in pediatric cohorts.

Age-localized molecular “waves” have been described in proteomic and metabolomic studies, particularly around adult transition periods^10,11,20^. Our analysis extends this transition-window concept to cell-type-resolved immune transcriptomes and further shows that the dominant contributing lineages shift across stages of life. Early-childhood remodeling was dominated by monocyte and B-lineage compartments, whereas T-cell subsets ranked more prominently in later transition windows, placing the widely reported age sensitivity of T-cell states^7,8^ into a stage-dependent context. This window-by-lineage perspective refines the interpretation of age effects in immune compartments and helps prioritize specific cell types and time periods for mechanistic follow-up.

Immunometabolic remodeling is well established as an important contributor to immune-cell function and plasticity in adults^3^, whereas the timing and cell-type breadth of immune-cell metabolic programs during childhood remain less well characterized. Our analysis identified a prominent early-childhood immunometabolic transition centered around age 6, with remodeling of metabolism-related transcriptional programs detected across multiple immune cell subsets. Within this transition window, CD16⁺ monocytes exhibited significant overrepresentation of the oxidative phosphorylation–related genes among window-associated DEGs, consistent with recent evidence implicating oxidative phosphorylation as a key feature of early-life monocyte immunometabolism associated with myeloid differentiation and inflammatory responsiveness^2^. This transition window also showed enrichment of genes annotated to glutathione and pyruvate metabolism in naïve B cells, pathways implicated in B-cell activation and class-switch recombination^3,21^. Together, these findings suggest that the 6-year transition captures a reorganization of metabolism-related immune transcriptional programs, potentially reflecting cell-type-specific metabolic adaptation accompanying immune-cell maturation and functional tuning during early childhood.

Early childhood also coincides with major endocrine–metabolic transitions beyond immune maturation. The lipidome undergoes extensive remodeling during the first 6 years of life^22^, dysregulation of which has been linked to immune-mediated diseases in childhood^23^. Also around age 6, adrenarche initiates a maturational rise in adrenal steroid secretion (including DHEA and related derivatives)^24,25^, and premature adrenarche has been associated with later low-grade inflammation^25,26^. Consistent with this developmental context, our serum profiling revealed a pronounced wave of lipid species and steroid-associated metabolites around ages 6-8. Notably, many of these wave-associated steroid derivatives and phospholipid species (particularly PC) showed significant associations with contemporaneous variation in immune-cell transcriptional programs and with features included in the transition scoring model that captures immune-state variation. Although cross-sectional co-variation cannot establish causality, these results highlight endocrine–metabolic signals as plausible correlates of immune-state remodeling during this interval.

Finally, we developed a transition score and an age-specific reference chart anchored to the 6-year immunometabolic transition in healthy individuals, which identified significant deviations across multiple pediatric immune disorders. These findings suggest that the biological programs captured by the score extend beyond normal age-associated variation and remain relevant in disease-related states. As an illustrative example, *SELL*, among the most influential genes in the scoring model, encodes L-selectin (CD62L), whose downregulation marks the naïve-to-effector transition^27^. Dysregulated CD62L expression has been reported in autoimmune conditions, including T1D^28^ and SLE^29^. The displacement of CD, SLE, T1D, and JIA from the healthy reference trajectory suggests the metabolism-related immune transcriptional programs established during this early childhood transition may be perturbed. Consistent with this interpretation, these diseases exhibit perturbations along the immune-metabolic axis, involving varying degrees of systemic metabolic alterations^30,31^ and immune-cell metabolic reprogramming^32–35^. Although whether the score can pinpoint specific immune functions or developmental processes remains to be determined, these deviations support the utility of this age-referenced framework for highlighting atypical immune-state patterns.

This study has several limitations. The cross-sectional design does not directly capture within-individual trajectories, and PBMC profiling cannot resolve potentially distinct tissue-resident immune programs. Moreover, metabolite– and lipid–transcript associations are observational and do not establish mechanistic relationships. Nevertheless, our study lays a foundation for mechanistic studies of the early-childhood immunometabolic transition and provides an age-resolved framework for interpreting pediatric immune variation.

## Methods

### Study design and cohorts

This study was designed to construct a lifespan-resolved single-cell atlas of peripheral immune cells using single-cell RNA sequencing (scRNA-seq) of peripheral blood mononuclear cells (PBMCs). We aggregated PBMC scRNA-seq data from 2,008 healthy donors spanning ages 0–100 years. Among these individuals, 1,952 donors were aggregated from 28 publicly available datasets (detailed dataset information is provided in Supplementary Table 1), with a median age of 55 years. We additionally recruited and profiled 56 healthy donors in-house.

We included PBMC scRNA-seq samples from individuals annotated as healthy, control, or normal in the original studies, with donor-level age and sex metadata available. All scRNA-seq data were generated using 10x Genomics platforms. For in-house recruitment, individuals were eligible if they were clinically healthy at the time of blood collection and had no history of chronic inflammatory or autoimmune disorders, hematologic diseases, or malignancy. Exclusion criteria included recent acute infection or fever, recent antibiotic use, and current use of systemic immunomodulatory or immunosuppressive medications. All samples collected by our group were obtained in accordance with the Declaration of Helsinki. Written informed consent was obtained from adult participants or, for minors, from a parent or legal guardian with assent when appropriate. The study was approved by the ethics committees of the First Affiliated Hospital of the University of Science and Technology of China (No. 2019KY027) and the First Affiliated Hospital of Bengbu Medical University (No. 2023KY017).

### Single-cell RNA-seq library preparation and sequencing

For samples generated by our group, frozen PBMCs were thawed and processed in 19 pooled batches using the 10x Genomics Chromium system according to the Chromium Single Cell 3′ Reagent Kits v3.1 protocol. Briefly, single cells were partitioned into gel bead–in-emulsions (GEMs) together with barcoded gel beads, enabling reverse transcription to generate cDNA molecules labeled with cell-specific barcodes and unique molecular identifiers (UMIs). The resulting cDNA was subsequently amplified by PCR to obtain sufficient material for library preparation. Libraries were sequenced on an Illumina NovaSeq 6000 platform with a target depth of at least 20,000 reads per cell. FASTQ files were then aligned to the GRCh38 reference genome and transcriptome using the CellRanger (v7.1.0) pipeline, which generated a gene-by-cell expression matrix. Donor demultiplexing and sample assignment were performed using Souporcell (v2.0)^36^.

### scRNA-seq quality control, preprocessing, and cell-type annotation

Scanpy (v1.10.3)^37^ was used to convert scRNA-seq data from mtx, txt, and csv formats into the standardized h5ad format. Samples were subsequently integrated by gene-wise intersection, retaining only genes consistently detected across all datasets. Raw count matrices were merged across samples using Scanpy (v1.10.3), followed by quality control filtering. Genes expressed in fewer than three cells were excluded, as were cells expressing fewer than 200 genes or more than 5,000 genes, as well as cells with mitochondrial gene content exceeding 25%. Putative doublets were identified and removed using DoubletDetection (v4.2)^38^ with parameters p_thresh = 1× 10^-16^ and voter_thresh = 0.5.

Cell identities were predicted using CellTypist (v1.6.3)^15^. The dataset comprising 1,265,624 PBMCs from Kock KH et al.^39^ was used to train a CellTypist classifier, which was subsequently applied for annotation prediction. Cell type annotations were inferred using celltypist.annotate with default parameters. Annotation accuracy was evaluated based on the CellTypist confidence scores and the expression level of canonical marker genes. Platelets and hematopoietic stem and progenitor cells were removed from downstream analyses.

### Lifespan dynamics of immune cell composition

To quantify age-associated changes in PBMC composition and assess their reproducibility across cohorts, we modeled the donor-level proportion of each immune cell subset as a smooth function of chronological age using generalized additive models (GAMs) and evaluated on a 300-point age grid. For each of the 26 cell types in the discovery cohort, we fitted the following model:

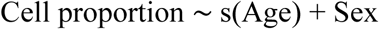

using restricted maximum likelihood (REML) with shrinkage/selection (select = TRUE) and γ = 1.2, with spline basis dimension *k* = 10. Sex-marginalized trajectories were derived by generating sex-specific predictions and averaging them using the cohort’s empirical sex distribution; predicted trajectories were then standardized as z-scores across the age grid.

### Pseudobulk gene expression preprocessing

For each donor and immune cell subset, raw counts were aggregated across cells to generate pseudobulk gene expression profiles. Genes expressed in fewer than 100 individuals were removed, followed by normalization using the trimmed mean of M values (TMM) method in the R package edgeR (v4.4.0)^40^. Normalized expression values were converted to log-transformed counts per million for downstream analyses. Lme4 (v1.1-35.5)^41^ was used to account for potential confounding effects for each gene. Age and sex were included as fixed effects, and data source was included as a random intercept. Adjusted expression values were obtained by removing the effects of sex and data source while retaining the fitted age effect and residual variation. The adjusted gene expression matrices were used for subsequent analyses, including the identification of age-associated genes, differential expression-sliding window analysis (DE-SWAN), metabolite–transcript and lipid–transcript association analyses, construction of the transition scoring model and calculation of individual transition scores.

### Identification of linear and nonlinear age-associated genes

We identified genes exhibiting linear changes with chronological age by calculating Pearson correlation coefficients between gene expression levels and age using cor.test in R. Multiple testing correction was performed independently within each cell type using the Benjamini–Hochberg procedure to control the false discovery rate. Genes that met an adjusted P value threshold of < 0.05 were considered to show a significant linear association with age. In addition, genes with absolute Pearson correlation coefficients < 0.2 were excluded, as correlation coefficients below 0.20 are generally regarded as very weak or negligible^42^. Genes below this correlation threshold were interpreted as having only a limited proportion of their age-related variation explained by a linear model.

TradeSeq (v1.20.0) was used to identify genes from transcriptomic data that were differentially expressed along an ordered temporal trajectory. This approach applies GAMs, which can capture nonlinear associations^43^. For each cell type, chronological age and scaled gene expression values were used as inputs, with a Gaussian family specified for model fitting. P values were adjusted for multiple comparisons using the Benjamini–Hochberg method, and genes with an adjusted p value < 0.05 were considered significantly associated with age. Genes showing significant linear associations within each cell type were subsequently excluded, yielding a set of genes exhibiting nonlinear age-associated expression patterns.

The locally estimated scatterplot smoothing (LOESS) approach was applied to visualize the trajectories of gene expression across the lifespan. For each gene in each cell type, expression values were standardized as z-scores, and a LOESS regression model was fitted. During the fitting process, the LOESS span parameter was optimized by fivefold cross-validation over values ranging from 0.2 to 1.0. This ensured that the LOESS model provided an accurate and non-overfitting fit to the gene expression data^11^. Then the LOESS prediction models were applied to predict the expression of each gene at every 1-year age interval. For visualizing the overall transcriptional trajectories of immune cell subsets, fitted expression values across ages 0–100 years were z-scored for each gene within each cell subset, and the mean z-score across all genes was calculated at each age point to represent the aggregate transcriptional state.

### Quantifying waves of transcriptomic change

The DE-SWAN algorithm^10^ was used to quantify localized waves of transcriptomic change across the lifespan, leveraging the sex- and data source-adjusted gene expression matrices generated as described above. A sliding window with a predefined age span was moved along the continuous age axis at a fixed step size. At each window center, individuals within the window were divided into younger and older groups relative to the center age, and gene expression levels were compared between the two groups using the following model:

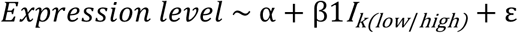

*I_k_*_(*low/high*)_ represents a binary indicator of whether an individual was younger or older than the center age (*k*) within the corresponding window. Differentially expressed genes (DEGs) were identified independently at each window center. P values were adjusted for multiple testing using the Benjamini–Hochberg method, and genes with an adjusted P value < 0.05 were considered significant. Based on the age distribution of the samples and preliminary analyses, the sliding-window width was set to 8 years and advanced in 2-year steps. For each cell type, peaks of transcriptomic waves were defined as window-centered ages at which the number of DEGs exceeded that at the immediately adjacent age centers.

### Functional enrichment analyses

Pathway enrichment analyses based on the GO and KEGG databases were performed using Metascape for Bioinformaticians (MSBio, v3.5)^44^ with the parameters min_overlap = 3 and min_enrichment = 1.5. To further filter and categorize metabolic pathways from the KEGG enrichment results, pathway annotations were curated by referring to the KEGG database and the classification scheme described by Zhou et al.^45^, resulting in the organization of KEGG pathways into 10 major metabolic categories.

### Serum metabolomics and lipidomics profiling

Ultra-performance liquid chromatography–mass spectrometry (UPLC–MS) analysis and preprocessing of serum metabolomics data

Metabolites were extracted using a ternary solvent system consisting of methanol, acetonitrile, and water in a 4:2:1 ratio. Extracts were dried and subsequently reconstituted in a 1:1 (v/v) methanol–water solution prior to analysis. Quality control samples were prepared by pooling 20 μL from each individual sample. Untargeted metabolomic profiling was performed using a UPLC–MS platform operated in both positive and negative ion modes. Chromatographic separation was performed on a BEH C18 column (1.7 μm, 2.1 × 100 mm; Waters, USA). In positive ion mode, mobile phase A consisted of 0.1% formic acid in water, and mobile phase B consisted of acetonitrile. In negative ion mode, mobile phase A consisted of 10 mM ammonium formate in water, and mobile phase B consisted of acetonitrile. Data acquisition was carried out on a Thermo Q Exactive mass spectrometer (Thermo Fisher Scientific, USA) with a scan range of m/z 70–1,050. Raw mass spectrometry data were processed using Compound Discoverer software (v3.3; Thermo Fisher Scientific, USA). Metabolic feature detection and annotation were performed by matching accurate mass and MS/MS spectra against the BGI Metabolome Database (BMDB), mzCloud, and the ChemSpider online database, generating a data matrix containing metabolite peak areas and identification information.

The result files from Compound Discoverer were imported into MetaX for data preprocessing and downstream analysis. Data preprocessing included the following steps: normalization by probabilistic quotient normalization (PQN) to obtain relative peak areas; batch-effect correction using quality control-based robust LOESS signal correction; and removal of metabolites with a coefficient of variation greater than 30% based on relative peak areas in quality control samples. Metabolite annotation and classification were performed using the Human Metabolome Database (HMDB) and KEGG pathway databases, and unannotated molecules were removed from downstream analyses. “Super Class” and “Class” annotations from HMDB were used to classify metabolites at different levels of granularity for subsequent comparisons of metabolite counts across categories.

### UPLC–MS analysis and preprocessing of serum lipidomics data

Lipids were extracted using pre-cooled isopropanol. Quality control samples were prepared by pooling 10 μL from each individual sample. Untargeted lipidomic profiling was performed using a UPLC–MS platform operated in both positive and negative ion modes. Chromatographic separation was performed on a BEH C18 column (1.7 μm, 2.1 × 100 mm, Waters, USA). In positive ion mode, mobile phase A consisted of 60% acetonitrile in water, 10 mM ammonium formate, and 0.1% formic acid, and mobile phase B consisted of 90% isopropanol, 10% acetonitrile, 10 mM ammonium formate, and 0.1% formic acid. In negative ion mode, mobile phase A consisted of 60% acetonitrile in water and 10 mM ammonium formate, and mobile phase B consisted of 90% isopropanol, 10% acetonitrile and 10 mM ammonium formate. MS1 and MS/MS data were acquired using a Q Exactive mass spectrometer (Thermo Fisher Scientific, USA). The full scan range was m/z 70‒1,050, with a resolution of 70,000. Raw mass spectrometry data were processed using LipidSearch software (v.4.1; Thermo Fisher Scientific, USA).

The result files from LipidSearch were imported into MetaX for data preprocessing and further analysis. Data preprocessing included the following steps: removal of lipids with more than 50% missing values in quality control samples or more than 80% missing values in experimental samples; imputation of missing values using the k-nearest neighbors algorithm; normalization by PQN to obtain relative peak areas; batch-effect correction using quality control-based robust LOESS signal correction; and removal of lipids with a coefficient of variation greater than 30% based on relative peak areas in quality control samples.

### Metabolomics and lipidomics analysis

Based on the age distribution of the samples and preliminary analyses, the DE-SWAN analyses for metabolomics and lipidomics were performed using a 4-year window, with the window sliding every 2 years from ages 4 to 20. P values were adjusted for multiple comparisons using the Benjamini–Hochberg method. The narrower window was selected because the pediatric metabolomics cohort contained a restricted age range. Significantly changed molecules were identified with thresholds of 0.05 for the adjusted P value.

For each immune cell type, Spearman correlation analyses were performed between the expression levels of DEGs and the abundances of serum molecules that were significantly altered in the corresponding transition age window (the 6–8-year window for metabolomics and the 6-year window for lipidomics). Correlation coefficients and P values were calculated using the rcorr function in the Hmisc R package (v5.2). P values were adjusted for multiple testing across all tested metabolite–transcript or lipid–transcript pairs using the Benjamini–Hochberg method. Significant correlations were defined by an adjusted P value < 0.05 and an absolute correlation coefficient > 0.2.

For significant metabolite–transcript and lipid–transcript associations, genes linked to metabolites annotated as Steroids and steroid derivatives or Carboxylic acids and derivatives in the HMDB Class, as well as genes associated with lysophosphatidylcholine (LPC) and phosphatidylcholine (PC) lipid species, were subjected to Gene Ontology (GO) Biological Process enrichment analysis. For each cell type, the five most significantly enriched pathways ranked by −log10(P value) were retained.

### External validation cohorts

To identify external validation cohorts for evaluating the reference chart, we searched PubMed for publicly available human peripheral blood single-cell RNA sequencing (scRNA-seq) datasets published up to April 24, 2026. The search strategy combined terms related to single-cell transcriptomics (“single-cell RNA sequencing”, “scRNA-sequencing”, or “single cell”) with terms related to peripheral blood mononuclear cells (“peripheral blood mononuclear cells” or “PBMCs”) and disease-specific terms for immune-related disorders. Eligible datasets were required to meet the following criteria: (1) generated using 10× Genomics scRNA-seq platforms; (2) derived from fresh human PBMC samples; and (3) accompanied by participant-level demographic information, including age and sex. For pediatric immune-related disease cohorts, only studies including participants younger than 21 years of age were considered eligible. After screening and eligibility assessment, we identified one healthy validation cohort and four pediatric immune-related disease cohorts for independent evaluation of the immune-state reference chart.

The healthy cohort (n=61)^7^ and three pediatric immune-related disease cohorts, including Crohn’s disease (CD; n = 7)^46^, systemic lupus erythematosus (SLE; n = 30)^47^, and Juvenile idiopathic arthritis (JIA; n = 6)^48^ were obtained from publicly available resources, whereas the pediatric type 1 diabetes (T1D; n = 109) cohort was generated in-house by our group.

T1D cases were included based on a clinical diagnosis of insulin-dependent diabetes with insulin treatment initiated at the time of diagnosis, together with additional evidence supporting this classification. Autoantibody positivity (GADA, IA-2A, or ZnT8A) was considered direct evidence supporting autoimmune diabetes. Participants were required to have a disease duration of less than 5 years at sample collection to minimize the potential impact of long-term disease progression and treatment exposure on immune-state profiles. All samples collected by our group were obtained in accordance with the Declaration of Helsinki. Written informed consent was obtained from a parent or legal guardian, with assent obtained when appropriate. The study was approved by the ethics committees of the First Affiliated Hospital of the University of Science and Technology of China (No. 2019KY027) and the First Affiliated Hospital of Bengbu Medical University (No. 2023KY017).

PBMC scRNA-seq data from all external validation cohorts were processed using the same pipeline for quality control, preprocessing, cell-type annotation, pseudobulk gene expression preprocessing, and adjustment for sex and batch, as described above. Individuals with missing values for selected features were removed. The resulting cell-type-specific pseudobulk gene expression matrices were used to extract the expression values of the 140 model features for each individual, which were then input into the transition scoring model, as described below, to calculate individual transition scores.

### Construction of the transition score

The discovery cohort, consisting of 2,008 healthy individuals used in the analyses described above, was used as the training cohort in this section. The transition scoring model was constructed using cell-type-specific DEGs identified within the 6-year transition window across immune cell subsets as candidate features. Feature selection was performed in two steps. First, DEGs with significant metabolite–transcript or lipid–transcript associations were retained. Second, recursive feature elimination with cross-validation (RFECV), implemented in scikit-learn (v1.7.2), was applied for further feature selection in the training cohort with cv = 5.

Using these selected features, a transition model was trained to classify individuals into pre-6-year (age ≤ 6) versus post-6-year (age > 6) states. Given the class imbalance between these two groups, model training was performed using the Self-paced Ensemble classifier (implemented in IMBENS v0.2.3), a method designed for class-imbalanced learning^49^, with logistic regression (C = 0.1) as the base estimator. The predicted probability output from the model was defined as the transition score, ranging from 0 to 1, with values closer to 0 indicating a pre-6-year transition state and values closer to 1 indicating a post-6-year transition state. Feature selection and model training were performed exclusively in the training cohort.

Feature importance was evaluated on the healthy validation cohort using the permutation_importance function in scikit-learn (v1.7.2), with n_repeats = 100. Relative importance was computed by dividing each feature’s importance by the maximum importance across all features, such that a value of 1 indicated the most important feature.

### Construction of the age-specific reference chart and evaluation of immune-state deviations

Transition scores from the training cohort were subsequently used to construct the age-specific reference chart using generalized additive models for location, scale, and shape (GAMLSS), a robust and flexible framework widely used for modelling nonlinear growth trajectories, including the World Health Organization growth standards^50^. Multiple transformations, including the Box-Cox power exponential (BCPE), Box-Cox t (BCT), and Box-Cox Cole and Green (BCCG) models^51^, were evaluated based on Akaike information criterion (AIC) and Bayesian information criterion (BIC). Although the BCPE model showed a slightly lower AIC, the BCCG model achieved the lowest BIC and provided a more parsimonious fit with fewer degrees of freedom. Therefore, the BCCG distribution was selected for subsequent reference curve construction (Supplementary Table 15). Model fit was further evaluated using visual inspection of residual distributions, worm plots, and Q-statistics. All analyses were performed using the gamlss package (v5.5-0) in R.

Using the fitted BCCG model, individualized z-scores were calculated for the training cohort, the healthy validation cohort, and pediatric cohorts with immune disorders. For each individual, age-specific μ, σ, and ν parameters were obtained from the reference model. The observed transition score was transformed into an age-standardized z-score using the Box-Cox transformation of the BCCG distribution, where μ, σ, and ν define the age-dependent location, scale, and skewness of the reference distribution, respectively.

Differences between the healthy training reference population and the independent healthy validation cohort or pediatric immune-related disease cohorts were assessed using permutation testing with 10,000 permutations. In each comparison, the observed difference in median z-scores between groups was compared against a null distribution generated by random label permutation. Two-sided empirical P values were calculated as the proportion of permuted differences greater than or equal to the observed difference in absolute value. Multiple-testing correction was performed using the Benjamini–Hochberg procedure.

## Acknowledgements

We thank the USTC Supercomputing Center and the School of Life Science Bioinformatics Center for providing computing resources for this project. This work was supported by the Noncommunicable Chronic Diseases-National Science and Technology Major Project (2023ZD0509100 to J.W), the National Natural Science Foundation of China grants (32570791, 32270978 to C.G., and 82470870 to X.Z.), and the Fundamental Research Funds for the Central Universities (WK9100000086 to C.G.).

## Author Contributions

J.W., C.G., and X.Z. conceived the study, secured funding, and supervised the overall research process; X.Z. contacted clinical samples and interpreted clinical data with the help of H.T. and T.Y.; J.Z. and Y.Shi. designed study, curated public data and conducted the study with the help of Y.Sun, Y.W., P.W. and R.L.; K.Q. provided consultation on the study design and manuscript writing; J.W., C.G., X.Z., J.Z., and Y.S. wrote the manuscript with the help of all the other authors. All authors contributed to the article and approved the submitted version.

## Data Availability

The in-house-generated scRNA-seq raw FASTQ files have been deposited in the Genome Sequence Archive for Human under accession no. HRA008320. The corresponding processed count matrices (h5ad) have been deposited in OMIX, China National Center for Bioinformation / Beijing Institute of Genomics, Chinese Academy of Sciences, under accession no. OMIX006887. The metabolomic and lipidomic data generated in this study have also been deposited in OMIX under accession no. OMIX012572.

Previously published scRNA-seq data included in this study, comprising raw FASTQ files or processed count matrices, were obtained from GEO, Synapse, Genome Sequence Archive, CELLxGENE Data Portal, and Zenodo. Details of all datasets are provided in Supplementary Table 1 and Supplementary Table 16.

## Code Availability

Code used for data analyses is available at https://github.com/JassicaZ/nonlinear-immune-remodeling.git. A reproducible, user-friendly pipeline for transition score calculation and reference chart visualization, along with a worked example, is also provided on GitHub at https://github.com/JassicaZ/6transition-scoring.

