## Extended Data Figure for "A lifespan-scale single-cell atlas defines an early-childhood immunometabolic transition linked to age-referenced immune states"

**Extended Data Figure Content**

Extended Data Fig. 1. Data integration and immune cell type annotation in the discovery cohort.

Extended Data Fig. 2. Age-associated genes with linear and nonlinear trajectories across cell types.

Extended Data Fig. 3. Distinct features of transcriptional remodeling across four transition windows.

Extended Data Fig. 4. Age-resolved metabolic pathway remodeling during childhood.

Extended Data Fig. 5. Lipidomic profiling for the pediatric subset.

Extended Data Fig. 6. Lipid–transcript associations in the pediatric subset.

Extended Data Fig. 7. Characterization of the transition scoring model.

Extended Data Fig. 8. Evaluation of the age-specific reference chart.

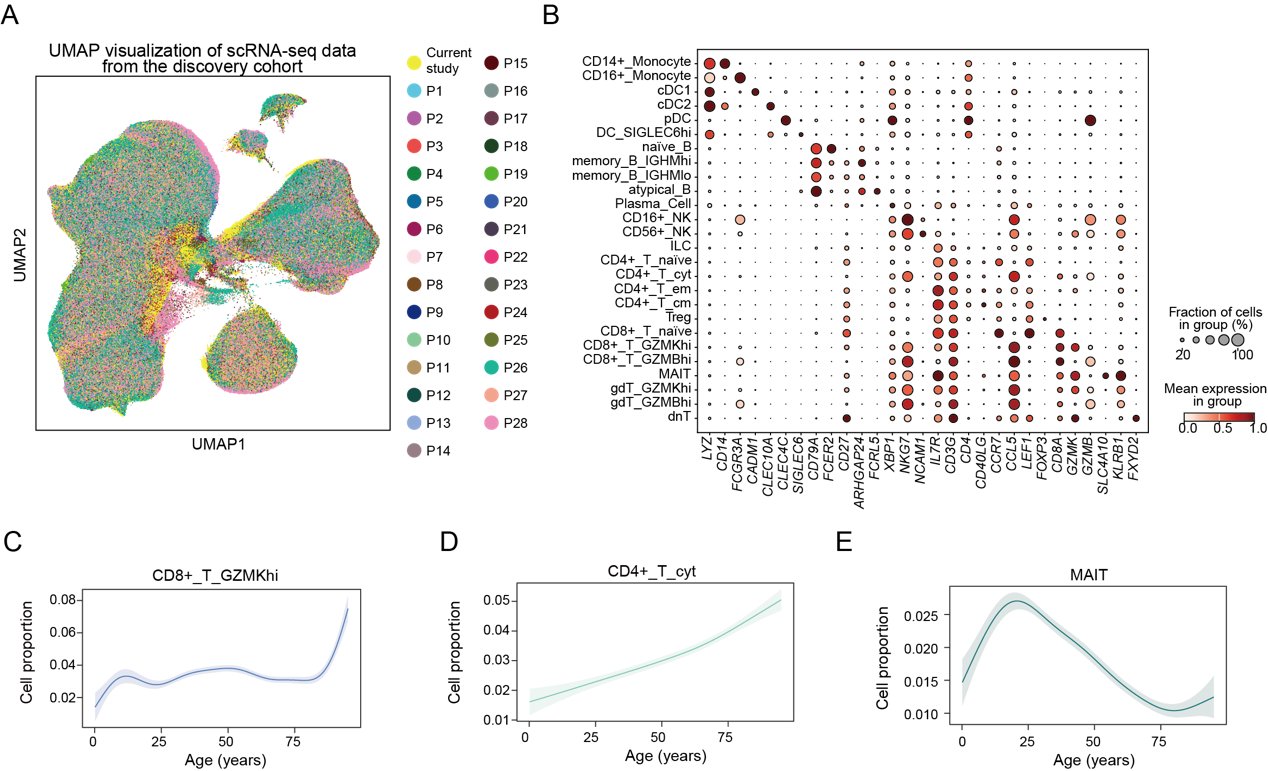

**Extended Data Fig. 1: Data integration and immune cell type annotation in the discovery cohort.** **A,** Uniform manifold approximation and projection (UMAP) visualization of peripheral blood mononuclear cell (PBMC) single-cell transcriptomes from the discovery cohort, colored by data source. **B,** Dot plot showing the expression of canonical marker genes in each cell type. Dot size indicates the percentage of cells expressing each marker gene within each cell type, and color intensity indicates the mean expression level. **C–E,** Representative age-associated trajectories of immune cell subset proportions across the human lifespan. Lines represent fitted smooth trajectories, and shaded areas indicate 95% confidence intervals.

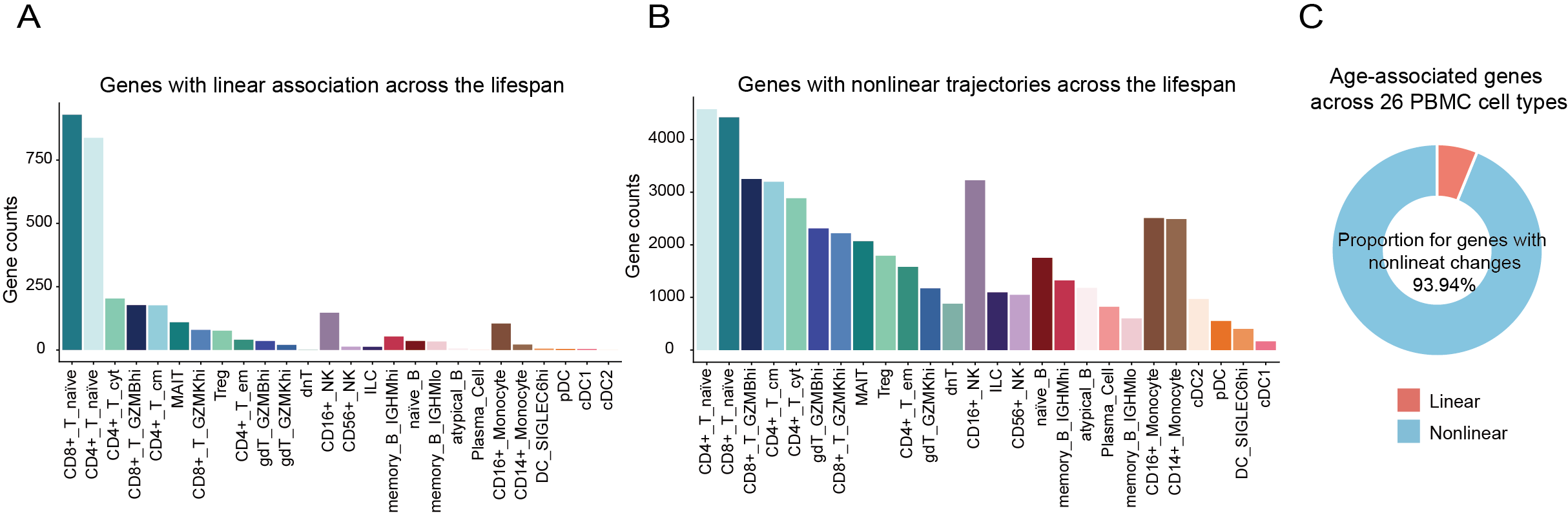

**Extended Data Fig. 2: Age-associated genes with linear and nonlinear trajectories across cell types. A,** Bar plot showing the number of genes with significant linear associations with age in each cell type. **B,** Bar plot showing the number of genes exhibiting nonlinear trajectories with age in each cell type. **C,** Pie plot showing the proportion of genes with linear and nonlinear associations with age. Genes exhibiting nonlinear age-related changes comprised the majority of age-associated genes.

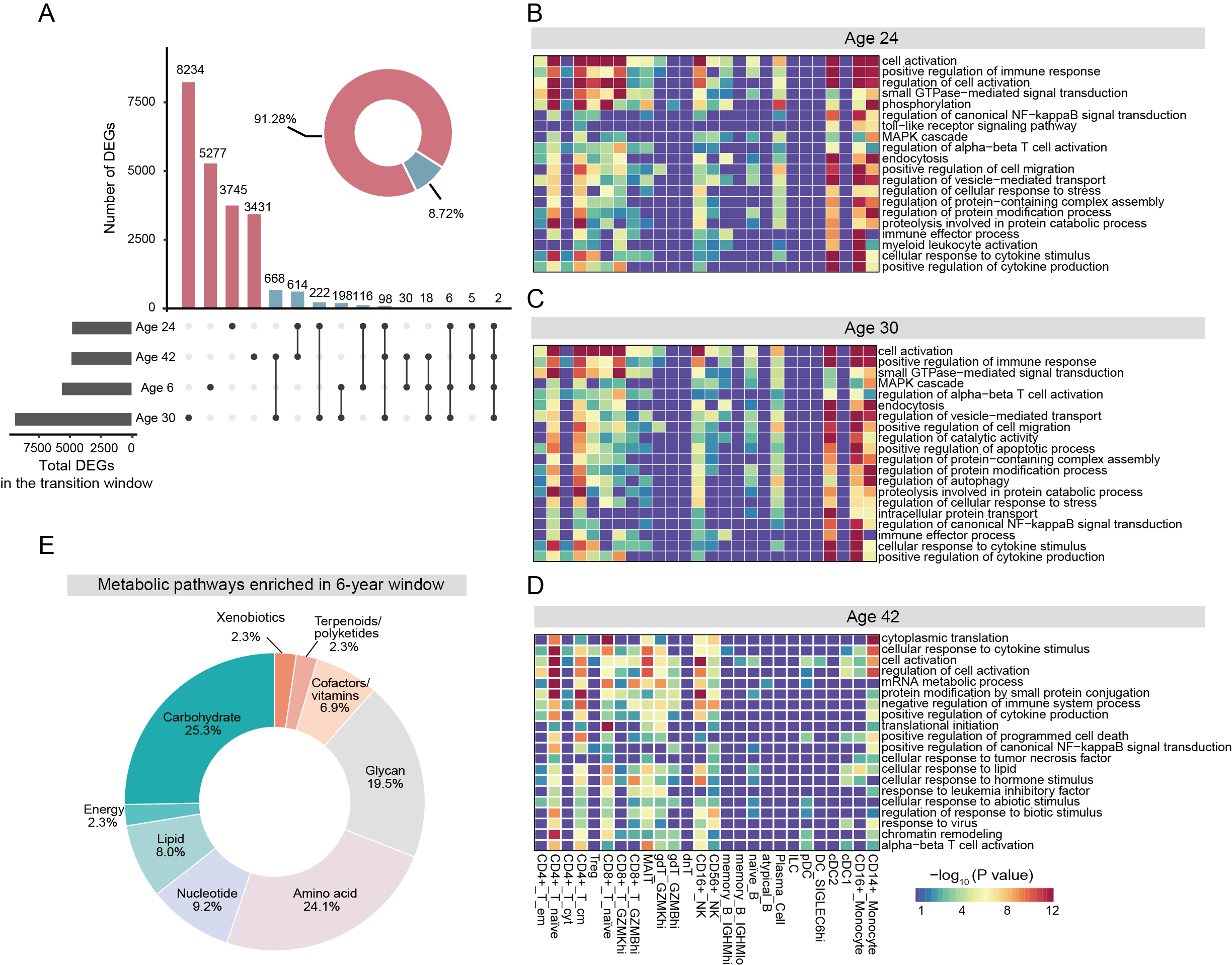

**Extended Data Fig. 3: Distinct features of transcriptional remodeling across four transition windows. A,** Overview of transcriptional remodeling across the four transition windows centered at 6, 24, 30, and 42 years. The upset plot shows the number of overlapping and window-specific differentially expressed genes (DEGs) across the four transition windows. The horizontal bar plot on the left shows the total number of DEGs detected in each transition window, while the upper bar plot indicates the number of DEGs shared across different combinations of transition windows. The pie chart in the upper right indicates the proportion of window-specific DEGs (pink) and shared DEGs (blue). **B–D,** Heatmaps illustrating the enrichment of biological processes among DEGs in 24, 30, and 42-year transition windows across immune cell subsets. **E,** Pie chart showing the distribution of significantly enriched KEGG metabolic pathway categories among DEGs from the 6-year transition window. Colors indicate different metabolic pathway categories.

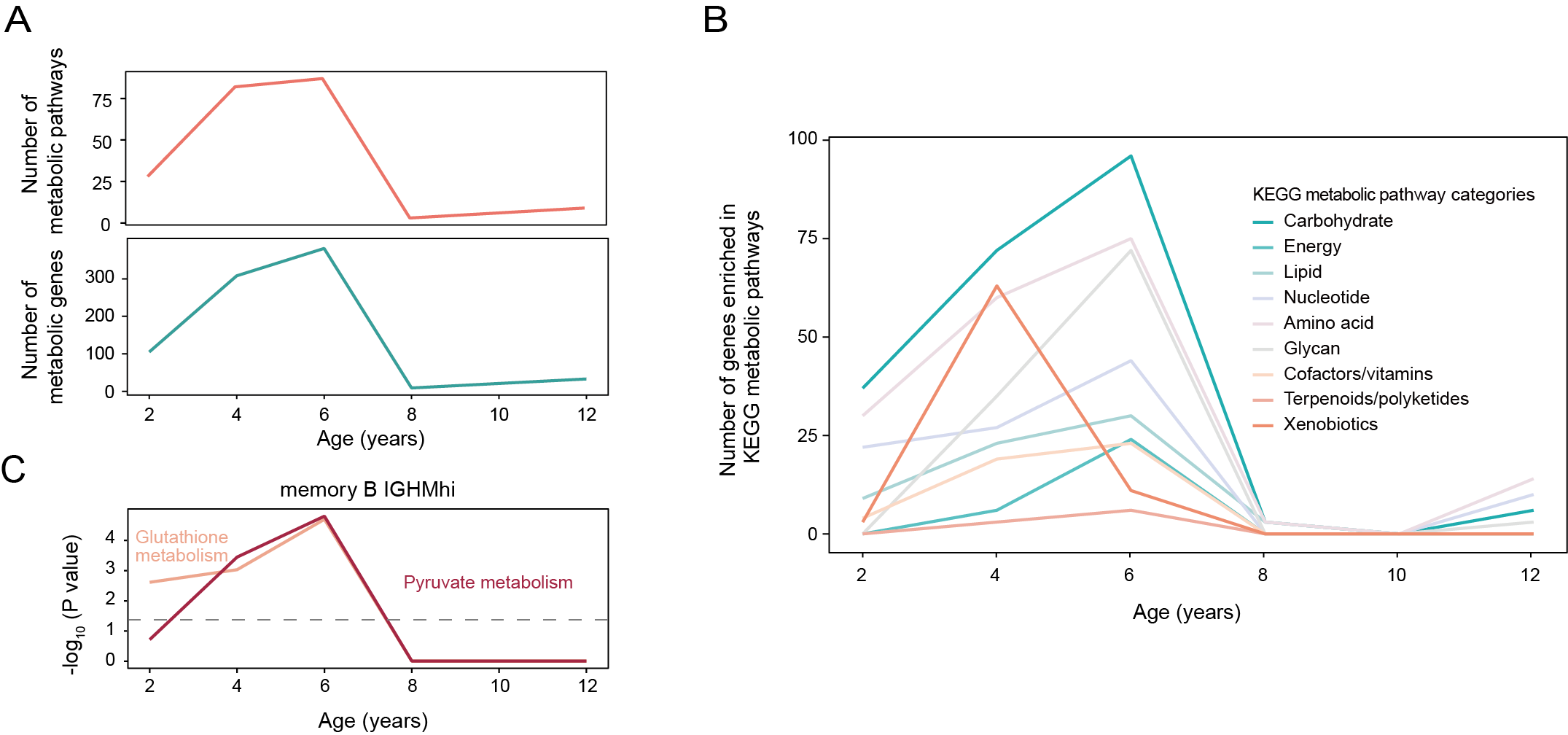

**Extended Data Fig. 4: Age-resolved metabolic pathway remodeling during childhood. A,** Line plots showing the number of significantly enriched metabolic pathways (top) and the number of genes involved in these pathways (bottom) across early childhood. **B,** Line plot depicting the number of genes contributing to enriched KEGG metabolic pathway categories. **C,** Line plots illustrating the significance of metabolic pathway enrichment, expressed as -log10(P value), for glutathione and pyruvate metabolism in IGHM^hi^ memory B cells. The dashed grey lines indicate P = 0.05.

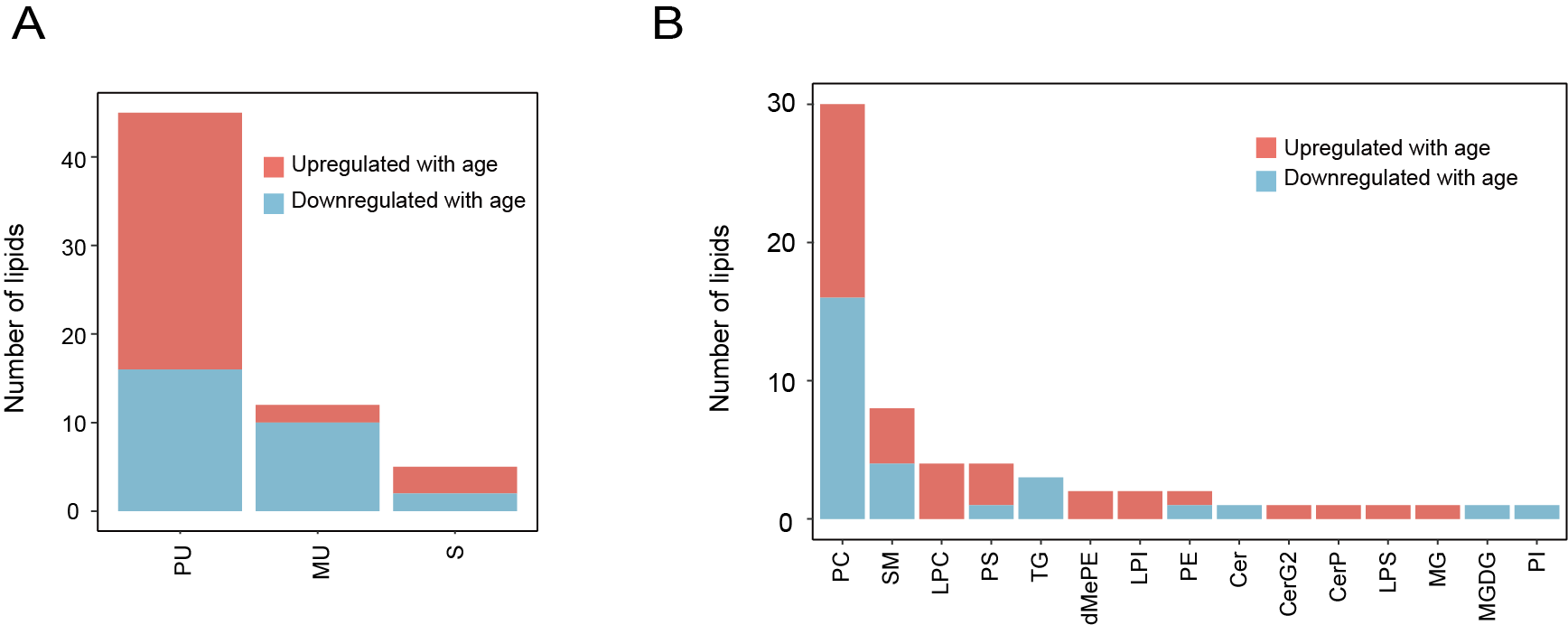

**Extended Data Fig. 5: Lipidomic profiling for the pediatric subset. A,** Stacked bar chart showing the number of lipids significantly upregulated (red) or downregulated (blue) with age within the 6-year transition window, categorized by degree of unsaturation. PU, polyunsaturated (≥ 2 double bonds); MU, monounsaturated (1 double bond); S, saturated (0 double bonds). **B,** Stacked bar chart showing the number of lipids significantly upregulated (red) or downregulated (blue) with age within the 6-year transition window across lipid classes. Cer, ceramides; CerG2, diglycosylceramide; CerP, ceramides phosphate; dMePE, dimethyl phosphatidylethanolamine; LPC, lyso-phosphatidylcholine; LPI, lyso-phosphatidylinositol; LPS, lyso-phosphatidylserine; MG, monoglyceride; MGDG, monogalactosyldiacylglycerol; PC, phosphatidylcholine; PE, phosphatidylethanolamine; PI, phosphatidylinositol; PS, phosphatidylserine; SM, sphingomyelin; TG, Triglyceride.

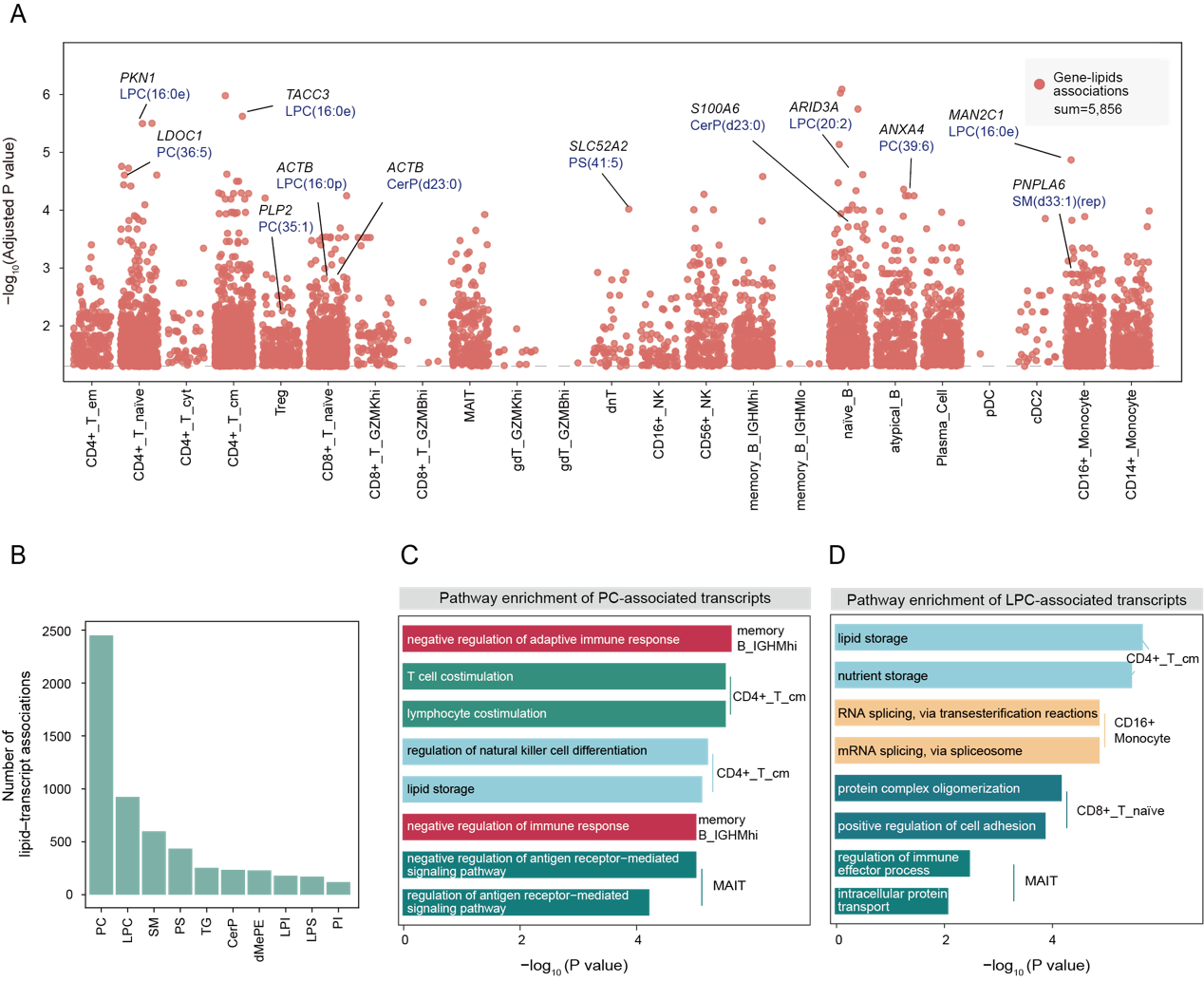

**Extended Data Fig. 6: Lipid–transcript associations in the pediatric subset. A,** Dot plot showing the significant lipid–transcript associations across immune cell subsets. The dashed grey lines indicate P = 0.05. **B,** Bar plot showing the number of lipid–transcript associations across different lipid species. **C**–**D,** Bar plots showing the functional enrichment of the age-localized DEGs associated with PC (C) and LPC (D). Bars indicate enriched pathways ranked by −log10(P value), with colors corresponding to the immune cell subsets indicated on the right.

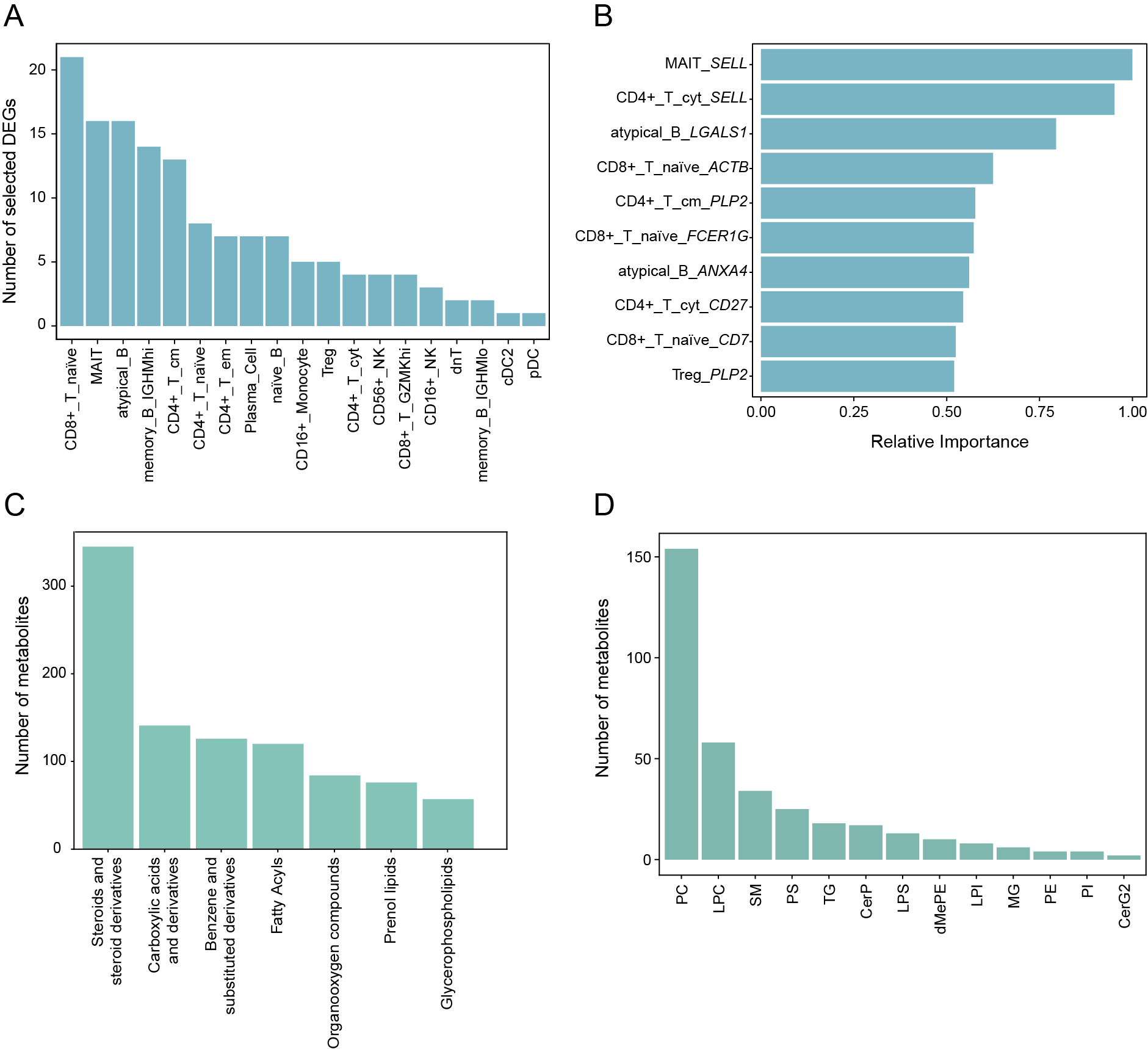

**Extended Data Fig. 7: Characterization of the transition scoring model. A,** Bar plot showing the number of 6-year transition-associated DEGs selected from each immune cell subset for inclusion in the transition scoring model. **B,** Bar plot showing the top 10 features ranked by relative importance in the scoring model. **C,** Bar plot showing the number of metabolite species associated with selected features. **D,** Bar plot showing the number of lipid species associated with selected features.

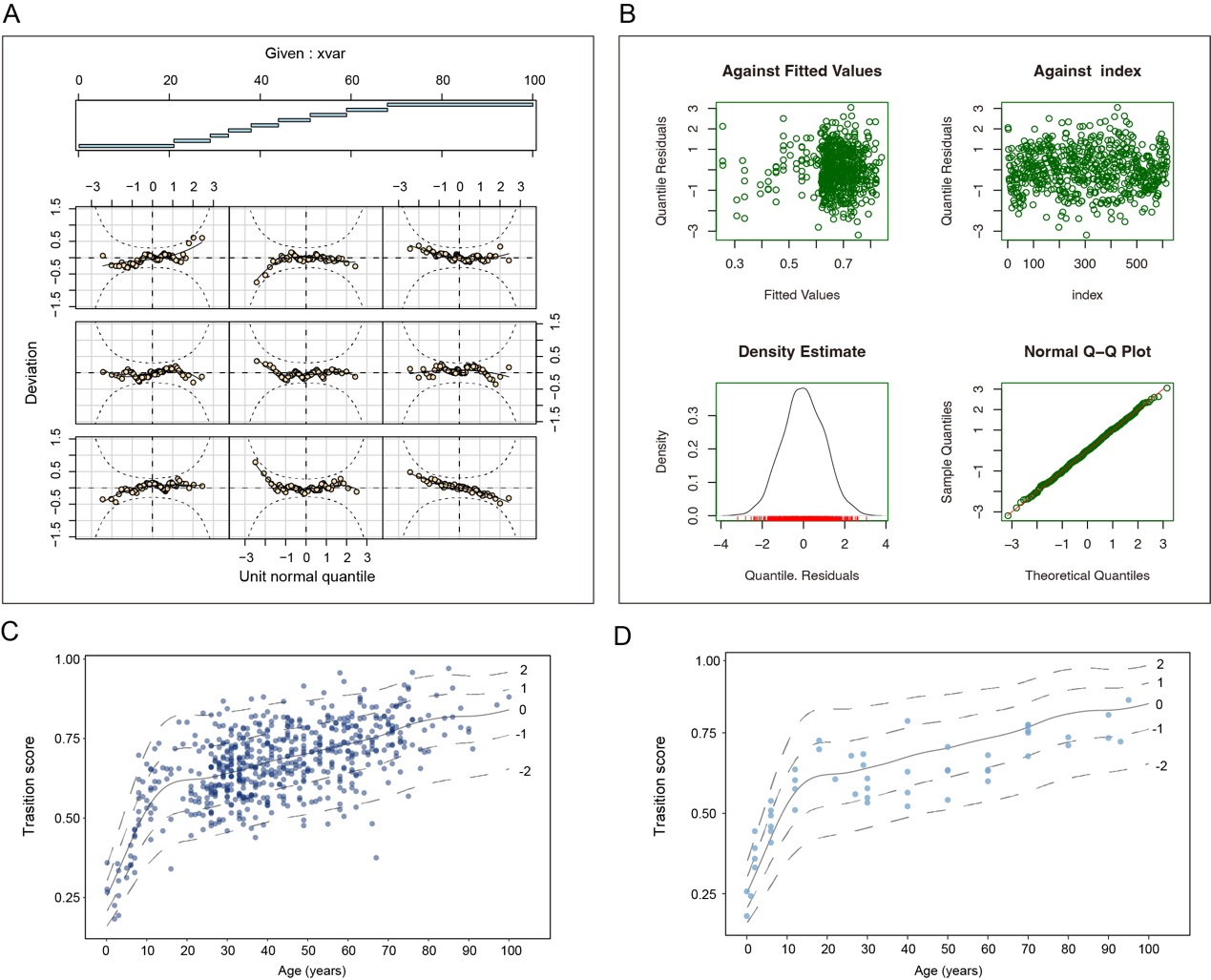

**Extended Data Fig. 8: Evaluation of the age-specific reference chart. A,** Worm plots evaluating the fit of the generalized additive models for location, scale and shape (GAMLSS). The upper panel shows the distribution of observations across age intervals used for worm plot construction. The lower panels display worm plots, presented as detrended Q**–**Q-plots of normalized quantile residuals within each age interval. **B, Residual diagnostic plots for the fitted GAMLSS model, including residuals plotted against fitted values (upper left), residuals plotted against observation index (upper right), a density estimate of quantile residuals (lower left), and a normal Q–Q plot of residuals (lower right).** **C–D,** Age-specific reference charts for the transition score. The solid line represents the age-specific reference median of healthy individuals (z-score = 0), and dashed lines indicate z-scores ranging from −2 to 2, from bottom to top. Points represent the individuals in the training cohort (C) and the healthy validation cohort (D).
